# *Drosophila* Kruppel homolog 1 represses lipolysis through interaction with dFOXO

**DOI:** 10.1101/165456

**Authors:** Ping Kang, Kai Chang, Ying Liu, Mark Bouska, Galina Karashchuk, Rachel Thakore, Wenjing Zheng, Stephanie Post, Colin S. Brent, Sheng Li, Marc Tatar, Hua Bai

## Abstract

Transcriptional coordination is a vital process contributing to metabolic homeostasis. As one of the key nodes in the metabolic network, the forkhead transcription factor FOXO has been shown to interact with diverse transcription co-factors and integrate signals from multiple pathways to control metabolism, oxidative stress response, and cell cycle. Recently, insulin/FOXO signaling has been implicated in the regulation of insect development via the interaction with insect hormones, such as ecdysone and juvenile hormone. In this study, we identified an interaction between dFOXO and the zinc finger transcription factor Kruppel homolog 1 (Kr-h1), one of the key players in juvenile hormone signaling in *Drosophila*. We found that *Kr-h1* mutants have reduced triglyceride storage, decreased insulin signaling and delayed larval development. Notably, Kr-h1 physically and genetically interacts with dFOXO *in vitro* and *in vivo* to regulate the transcriptional activation of adipose lipase *brummer* (*bmm*). The transcriptional co-regulation by Kr-h1 and dFOXO may represent a broad mechanism by which Kruppel-like factors integrate with insulin signaling to maintain metabolic homeostasis and coordinate organism growth.

## Introduction

Metabolic homeostasis plays important roles in developing animals ^1, 2^. The ability to coordinate growth and development with nutrient availability is critical for the adaptation to fluctuating environment. The main hormonal pathway that regulates insect growth and energy metabolism is insulin/insulin-like growth factor signaling (IIS). Unlike the single insulin, two insulin-like growth factor (IGF) system in mammals, insects have multiple insulin-like peptides ^3, 4^. The activation of insulin/insulin-like growth factor signaling stimulates two major kinase cascades: the PI3K/AKT pathway and MAPK/ERK pathways ^5^. In particular the O subclass of the forkhead transcription factors (FOXO) are substrates of PI3K/AKT. Decreased cellular IIS leads to de-phosphorylation and nuclear translocation of FOXO and the transcriptional activation of FOXO target genes ^6, 7^. Besides IIS, FOXO transcriptional activity is modulated by several other pathways (e.g. AMPK, JNK and SIRT) through post-translational modification (PTM) that modulate FOXO binding to DNA or its co-activators ^6, 7^.

FOXO plays a key role in mediating the cross-talk between insulin signaling and other insect hormones (e.g. juvenile hormone (JH) and ecdysteroids) to coordinate insect growth, development and metabolic homeostasis ^8-10^. Molting hormone ecdysone regulates developmental timing by inhibiting insulin signaling and promoting the nuclear localization of *Drosophila* forkhead transcription factor (dFOXO) ^8^. During the non-feeding pupation stages of *Bombyx* silkworm, 20-hydroxyecdysone (20E) induces lipolysis and promotes transcriptional activation of two adipose lipases via the regulation of FOXO ^11^. On the other hand, the link between JH and insulin signaling was first demonstrated in *Drosophila* where insulin receptor (InR) mutants were seen to reduced JH biosynthesis ^12^. Recent studies on size control further suggest that JH controls growth rate through *Drosophila* FOXO ^10^. Interestingly, JH also regulates lipid metabolism via the interactions with FOXO in Tsetse flies ^9^. Across these studies, it remains unclear how JH interacts with nutrient signaling and whether JH directly acts on FOXO-mediated transcriptional control.

FOXO interacts with a number of transcription factors within the nucleus to activate or inhibit transcription of target genes ^13^. The interactions between FOXO and its binding partners contribute to the transcriptional specificity of FOXO and pleotropic functions of insulin/FOXO signaling. For instance, mouse FOXO1 interacts with PGC-1α in liver to modulate insulin-mediated gluconeogenesis ^14^; mammalian FOXO1 binds to Smad2/3 in response to TGF-beta signaling and regulates cell proliferation ^15^; mammalian FOXO transcription factors (FOXO3A and FOXO4) interacts with beta-Catenin of Wnt signaling to modulate cellular oxidative response ^16^. In *Drosophila*, dFOXO interacts with bZIP transcription factor REPTOR of Mechanistic target of rapamycin (mTOR) signaling to regulate growth and energy homeostasis ^17^. Interestingly, recent studies found that FOXO interacts with Ultraspiracle (Usp), a co-factor of the ecdysone receptor, to regulate ecdysone biosynthesis and developmental timing in *Drosophila* ^18^. To date, factors of JH signaling have not been identified to directly interact with FOXO.

Kruppel-like homolog 1 (Kr-h1) is a key regulator of insect molting and metamorphosis and a major effector in JH signaling ^19-21^. JH strongly induces the transcription of *Kr-h1* via its receptor Methoprene-tolerant (Met) ^20-22^. During insect development, Kr-h1 functions as a transcriptional repressor on neurogenesis of mushroom body and photoreceptor maturation ^23, 24^. Kr-h1 belongs to Kruppel-like factors (KLFs) protein family, a group of conserved C2H2 type zinc finger transcription factors. Unlike mammalian KLFs that contain three zinc finger DNA binding domains, *Drosophila* Kr-h1 has eight zinc finger motifs ^25^. KLFs are also closely related to transcription factor Sp1 (specificity protein 1). At least seventeen KLFs are identified in mammals ^26^. Both KLFs and Sp1-like factors recognized GC-rich DNA elements or CACCC-box in the promoters of target genes ^26^. While KLFs and Sp1 can function as both transcription activator and repressor, the N-terminus of KLFs contains a consensus motif PXDL(S/T) that is thought to interact with transcriptional co-repressor CtBP (C-terminal binding protein) ^27, 28^.Some KLFs also interact with transcriptional co-activators to enhance transcriptional activities. For instance, KLF1 is acetylated through its interaction with co-activators p300 and CREB-binding protein (CBP), which leads to elevated induction of target gene beta-globin ^29^.

In this study, we identified an interaction between dFOXO and the zinc finger transcription factor Kr-h1. While characterizing a *Drosophila Kr-h1* mutant, we found that Kr-h1 controls lipid metabolism and insulin signaling. Kr-h1 physically interacts with dFOXO and represses the transcriptional activation of dFOXO target genes, including insulin receptor (*InR*) and triglyceride lipase (*bmm or brummer*). The present study suggests a mechanism by which Kruppel-like factor Kr-h1 integrates with insulin/dFOXO signaling to control lipid metabolism and coordinate organism growth.

## Results

### *Kr-h1* mutants delay larval development and have reduced triglyceride

Here we study the role of *Drosophila* Kruppel-like factor Kr-h1 in larval development and metabolic control using a P-element insertion line *Kr-h1[7]* (also known as *Kr-h1[k04411]*) ^19, 30^. The P-element insertion is located within exon 1 of the *Kr-h1α* isoform and is reported to interfere with the transcription of *Kr-h1* isoforms. *Kr-h1[7]* homozygous mutants are partially viable during embryonic and larval development ^30^. We backcrossed this *Kr-h1[7]* allele into a *yw*^*R*^ background for seven generations, producing a line where heterozygotes prolonged developmental time to pupariation (Fig. 1A), and homozygotes arrest at either second or third instar larval stage. *Kr-h1* mRNA is largely reduced in homozygous animals based on primers for the common region of all three isoforms (Fig. 1B). Using a newly generated rabbit anti-Kr-h1 antibody, three major bands were detected in larval samples from wild-type and *Kr-h1[7]* heterozygotes (Fig. 1C). Each of these bands was significantly reduced in homozygous animals, although a novel protein band with a distinct molecular weight (around 51 kDa) was observed (Fig. 1C). The identify of this protein band is unknown, but may correspond to the novel transcript observed previously by northern blot in *Kr-h1[7]* homozygous mutants ^19, 30^.

**Fig 1.**
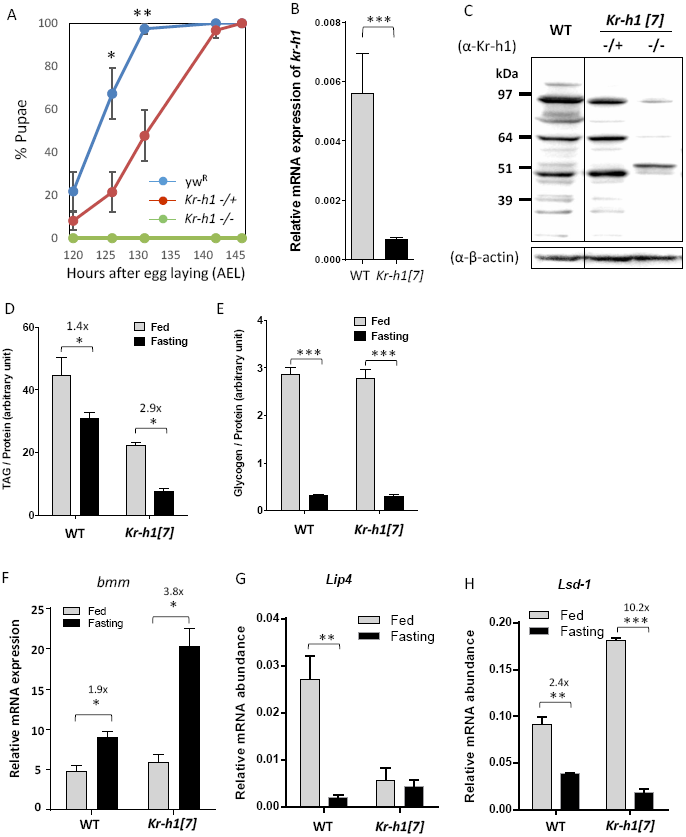
*Kr-h1* mutants delayed larval development and have reduced triglyceride. (A). *Kr-h1[7]* heterozygous mutants delayed pupation and homozygotes arrested at early larval stages. Percentage of pupariation at different developmental time points is shown. Data are represented as mean ± SE of three trials. Student t -test (** p<0.01, * p<0.05). (B). *Kr-h1* transcripts were significantly down-regulated in *Kr-h1[7]* mutants. Primers targeting common regions among three isoforms were used in qRT-PCR. Each bar represents mean ± SE of three biological replicates. Statistical significance between wild-type and mutants is assessed by student t-test (*** p<0.001). (C). Reduced Kr-h1 protein expression in *Kr-h1[7]* homozygous mutants. Larvae at 90 hr AEL (after egg laid) were used in western blots. In wild-type larvae, three distinct bands are found (∼84kDa, 64kDa and 48kDa). (D). *Kr-h1* mutant larvae have reduced TAG level. Upon starvation, TAG mobilization was faster in Kr-h1 mutants than in wild-type larvae. Larvae at 90 hr AEL were fasted for 16 hr in culture vials with wet kimwipe soaked with PBS. Each bar represents mean ± SE of three biological replicates. Statistical significance is assessed by two -way ANOVA followed by Tukey multiple comparisons test (*** p<0.001, ** p<0.01, * p<0.05). (E). Glycogen contents and the utilization rate were not affected by *Kr-h1* mutation. (F).Transcripts of TAG lipase *brummer* (*bmm*) were up-regulated by fasting and *Kr-h1* mutation. The fasting-induced *bmm* expression was further enhanced by *Kr-h1* mutation. (G). Transcripts of lysosomal acid lipase *lip4* were down-regulated by fasting and by *Kr-h1* mutation. (H). Fasting-triggered fly perilipin *Lsd-1* repression was significantly enhanced in *Kr-h1* mutants. Statistical significance is assessed by two-way ANOVA followed by Tukey multiple comparisons test (*** p<0.001, ** p<0.01, * p<0.05).

Defects in metabolic regulation also occur in the developmentally delayed *Kr-h1* mutants. Triglycerides and glycogen were measured in *Kr-h1[7]* homozygous larvae at 90 hours after egg laying (AEL). Among fed animals, triglycerides (TAG) were reduced 2-fold by *Kr-h1* mutation, while glycogen was similar among genotypes (Fig. 1D & 1E). TAG is a major stored nutrient mobilized during fasting. Accordingly, fasting reduced TAG stores in both genotypes, but to a significantly greater extent in *Kr-h1[7]* homozygous larvae (2.9-fold vs 1.4-fold in wild-type) (Two-way ANOVA, interaction p<0.047). Fasting reduced stored glycogen to the same extent in both genotypes (Fig. 1D & 1E).

In flies, adipose triglyceride lipase brummer (*bmm*) is a key lipase involved in TAG mobilization ^11, 31^. While transcripts of *bmm* were somewhat up-regulated by fasting in wildtype larvae and in fed *Kr-h1[7]* mutants (Fig. 1F), *bmm* expression was dramatically increased in fasted *Kr-h1[7]* homozygous larvae (4.3-fold vs. 1.8-fold in wildtype) (Two-way ANOVA, interaction p<0.0196) (Fig. 1F). This result is consistent with the greater TAG mobilization in fasted *Kr-h1* mutants as shown in Fig. 1D, suggesting that lipase activities might be enhanced in *Kr-h1* mutants, especially upon fasting. In parallel, we found that the expression of a lysosomal acid lipase *Lip4* was significantly down-regulated by fasting in wildtype and in fed *Kr-h1* mutants (Fig. 1G). These results suggest that Kr-h1 might specifically target the major adipose triglyceride lipase *bmm* to regulate TAG mobilization. As well, transcripts of fly perilipin *Lsd-1* were upregulated in *Kr-h1* mutants and down-regulated in both genotypes upon fasting (Fig. 1H). Perilipin proteins (PLINs) are a group of lipid droplet-associated proteins that act as protective coating factors to prevent lipid breakdown by triglyceride lipases ^32, 33^. Notably, repression of *Lsd-1* by fasting was significantly enhanced in *Kr-h1* mutants (10.2-fold vs. 2.4-fold in wildtype) (Two-way ANOVA, interaction p<0.0001). Collectively, these results suggest that Kr-h1 plays an important role in lipolysis through the transcriptional regulation of triglyceride lipase *bmm* and lipid droplet-associated protein *Lsd-1*.

### *Kr-h1* mutants have reduced insulin signaling

One way Kr-h1 might modulate TAG is through interactions with insulin/IGF signaling. Insulin/IGF signaling is a metabolic master regulator that controls lipase gene expression through its downstream transcription factor dFOXO ^34^. Here we see that phosphorylation of IIS-regulated kinase AKT was reduced in *Kr-h1[7]* homozygotes (Fig. 2A). Furthermore, *Kr-h1[7]* homozygotes had reduced expression of two insulin-like peptides (*dilp2* and *dilp5*), which are the major DILPs produced from brain neurosecretory cells, known as insulin producing cells (IPCs) (Fig. 2B & 2C).

**Fig 2.**
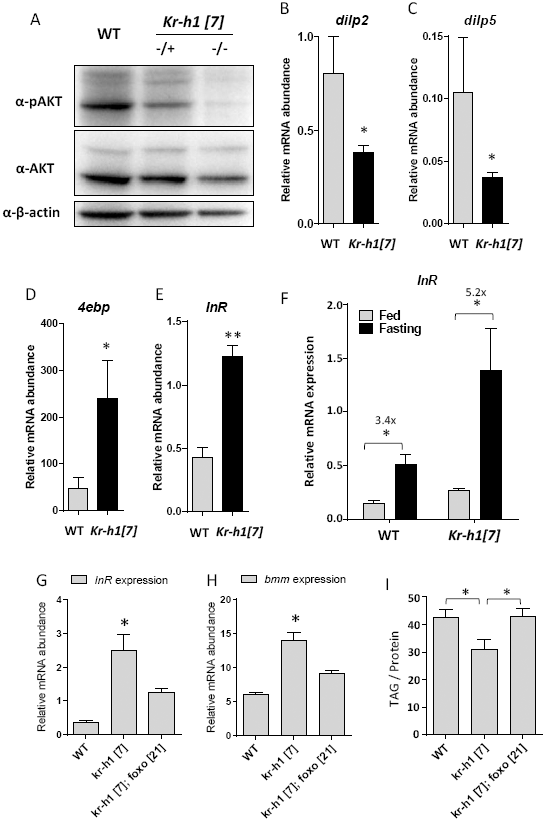
*Kr-h1* mutants have reduced insulin signaling. (A). Phosphorylation of AKT was down-regulated in *Kr-h1* mutants. Ten 90 hr AEL larvae were lysed in RIPA buffer and ∼20 μg of denatured protein was loaded to SDS-PAGE gels. (B-C). The transcripts of two insulin-like peptides (*dilp2*, *dilp5*) were down-regulated by *Kr-h1* mutation. (D-E).The mRNA expression of the key dFOXO targets *4ebp* and *InR* were up-regulated in *Kr-h1* mutants. Each bar represents mean ± SE of three biological replicates. Statistical significance between wild-type and mutants is assessed by student t-test (** p<0.01, * p<0.05). (F). *InR* transcripts is additively regulated by fasting and Kr-h1. Statistical significance is assessed by two-way ANOVA with Tukey multiple comparisons test (*** p<0.001, ** p<0.01, * p<0.05). (G). *dfoxo[21]* mutants suppress the induction of *InR* transcription by *Kr-h1[7]*. (H). *dfoxo[21]* mutants suppress the induction of *bmm* transcription by *Kr-h1[7]*. Each bar represents mean ± SE of three biological replicates. (I). *dfoxo[21]* mutants rescue the reduction of TAG levels in *Kr-h1[7]* mutants. Statistical significance is assessed by one-way ANOVA, followed by Dunnett’s multiple comparisons (* p<0.05).

Reduced insulin signaling is expected to activate forkhead transcription factor dFOXO ^6^. Accordingly, mRNA expression of two key dFOXO target genes, *4ebp* (eukaryotic translation initiation factor 4E binding protein) and *InR* were significantly induced in *Kr-h1* mutants (Fig. 2D & 2E), and *InR* expression was further increased in fasted *Kr-h1[7]* homozygotes (5.2-fold vs. 3.4-fold in wildtype) (Two-way ANOVA, interaction *p*=0.1023) (Fig. 2F). Thus, in *Kr-h1* mutant larvae, insulin signaling is inhibited and dFOXO is activated.

### *Kr-h1* genetically interacts with *dfoxo* to regulate the transcription of *InR* and *bmm*, and lipid metabolism

To determine the requirement of dFOXO for Kr-h1-mediated lipid metabolism, we generated a double mutant by combining *Kr-h1[7]* and *dfoxo[21]* ^35^. Interestingly, *dfoxo[21]* mutants suppressed the elevated *InR* and *bmm* expression found in *Kr-h1[7]* mutants (Fig. 2G & 2H), confirming that these transcription factors co-regulate key metabolic genes. Furthermore, the reduction of TAG in *Kr-h1[7]* mutants was rescued by *dfoxo[21]-/-* (Fig. 2I). Together, these results reveal a genetic interaction between Kr-h1 and dFOXO in the control of the transcription of metabolic genes and lipid metabolism.

### Kr-h1 physically interacts with dFOXO

Kr-h1 and dFOXO may interact directly or indirectly to regulate the expression of *InR* and *bmm*. To test the possibility of direct interaction, we attempted to co-immunoprecipitated (Co-IP) Kr-h1and dFOXO in cultured *Drosophila* cells. We were able to pull down endogenous dFOXO from nuclear and cytoplasmic extracts using an anti-dFOXO antibody. Interestingly, Kr-h1 was detected in the protein complex from the nuclear extracts, but not from the cytoplasmic extracts (Fig. 3A), suggesting that Kr-h1 can form a protein complex with dFOXO in the nuclei.

**Fig 3.**
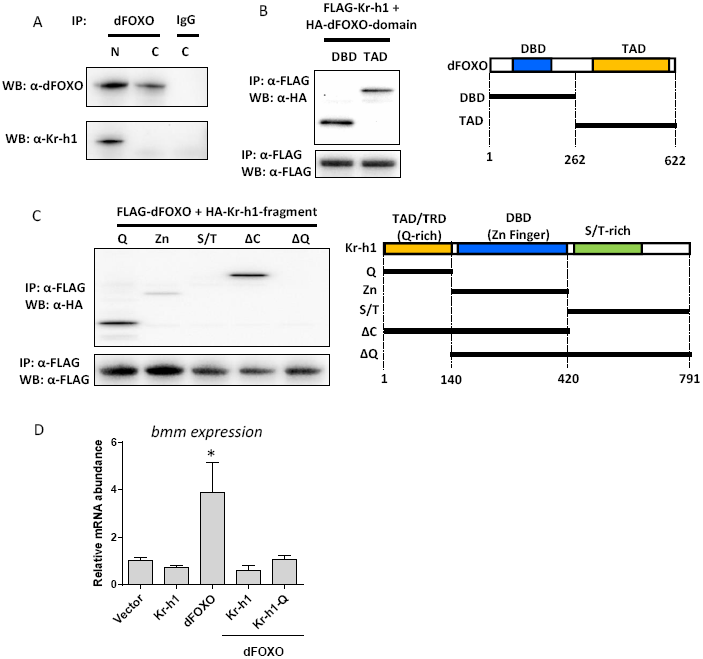
Kr-h1 physically interacts with dFOXO. (A). Co-immunoprecipitation of endogenous dFOXO and Kr-h1 from Kc167 cell lysates (N: Nuclear extracts; C: Cytoplasmic extracts). Anti-dFOXO antibodies were used in pull-down. Rabbit IgG served as a negative control. (B). Co-immunoprecipitation of FLAG-tagged full-length Kr-h1 and HA-tagged dFOXO fragments.Anti-FLAG antibodies were used to pull-down. Schematic graph on the right showing the position of each dFOXO fragment. Both DNA binding domain and transaction domain of dFOXO are able to bind to Kr-h1. (C). Co-immunoprecipitation of FLAG-tagged full-length dFOXO and HA-tagged Kr-h1 fragments. Anti-FLAG antibodies were used to pull down Kr-h1-dFOXO complex Schematic graph on the right showing the position of each Kr-h1 fragment. Q-rich domain shows strong binding to dFOXO. TAD/TRD: Transaction/repression domain. DBD: DNA binding domain. (D). Expression of either full-length Kr-h1 or Q-rich domain in Kc167 cells blocked dFOXO-induced transcription of *bmm*. Data are represented as mean ± SE of three trials. One-way ANOVA, followed by Dunnett’s multiple comparisons (* p<0.05).

To identify the protein interaction site between these transcriptional factors, we cloned a series of deletion fragments that contained different protein domains into the Gateway expression vectors. Both the DNA binding domain and transaction domain of dFOXO bound to full-length Kr-h1 proteins (Fig. 3B). On the other hand, the Kr-h1 fragments that contain transaction/repression domain (a Q-rich domain) bound to full-length dFOXO proteins, while Kr-h1 fragments with no Q-rich domain showed no binding (Fig. 3C). Therefore, the transaction/repression domain of Kr-h1 is responsible for the interaction between Kr-h1 and dFOXO.

The direct interaction between dFOXO and Kr-h1 may serve as a mechanism for the transcriptional repression of dFOXO target genes by Kr-h1. To test this idea, we co-expressed dFOXO with Kr-h1 (or Kr-h1 Q-rich domain) in Kc167 cells. The mRNA expression of *bmm* was significantly induced by dFOXO alone, and this induction was blocked by co-expressing either full-length of Kr-h1 or Q-rich domain (Fig. 3D). Thus, Kr-h1 appears to repress dFOXO transcriptional activity through direct protein-protein interactions.

### Kr-h1 binds to the promoters of *insulin receptor* and *brummer* lipase adjacent to dFOXO binding sites

Kr-h1 and dFOXO physically interact and may thus transcriptionally co-regulate metabolic genes. It has been previously shown that dFOXO binds to the promoter regions near transcriptional start sites of *InR* and *bmm* ^34, 36^, although our recent ChIP-Seq analysis (unpublished) suggests that dFOXO also strongly bound the promoter region near the 5’-UTR of *InR* (P1 region as shown in Fig. 4A) that contains a canonical FOXO binding motif (GTAAATAA). To identify potential Kr-h1 response elements of *InR* and *bmm*, we searched their promoters using mammalian KLF motifs in the Jaspar database (http://jaspar.genereg.net). Three putative KLF binding sites denoted P1∼P3 in each promoter were identified including sites in 5’-UTR and intronic regions (Fig. 4A & 4B) (Supplementary Table S1). We did not find any sites corresponding to the *Bombyx* Kr-h1 response element (GACCTACGCTAACGCTAAATAGAGTTCCGA) reported by Kayukawa et al. ^22^

**Fig 4.**
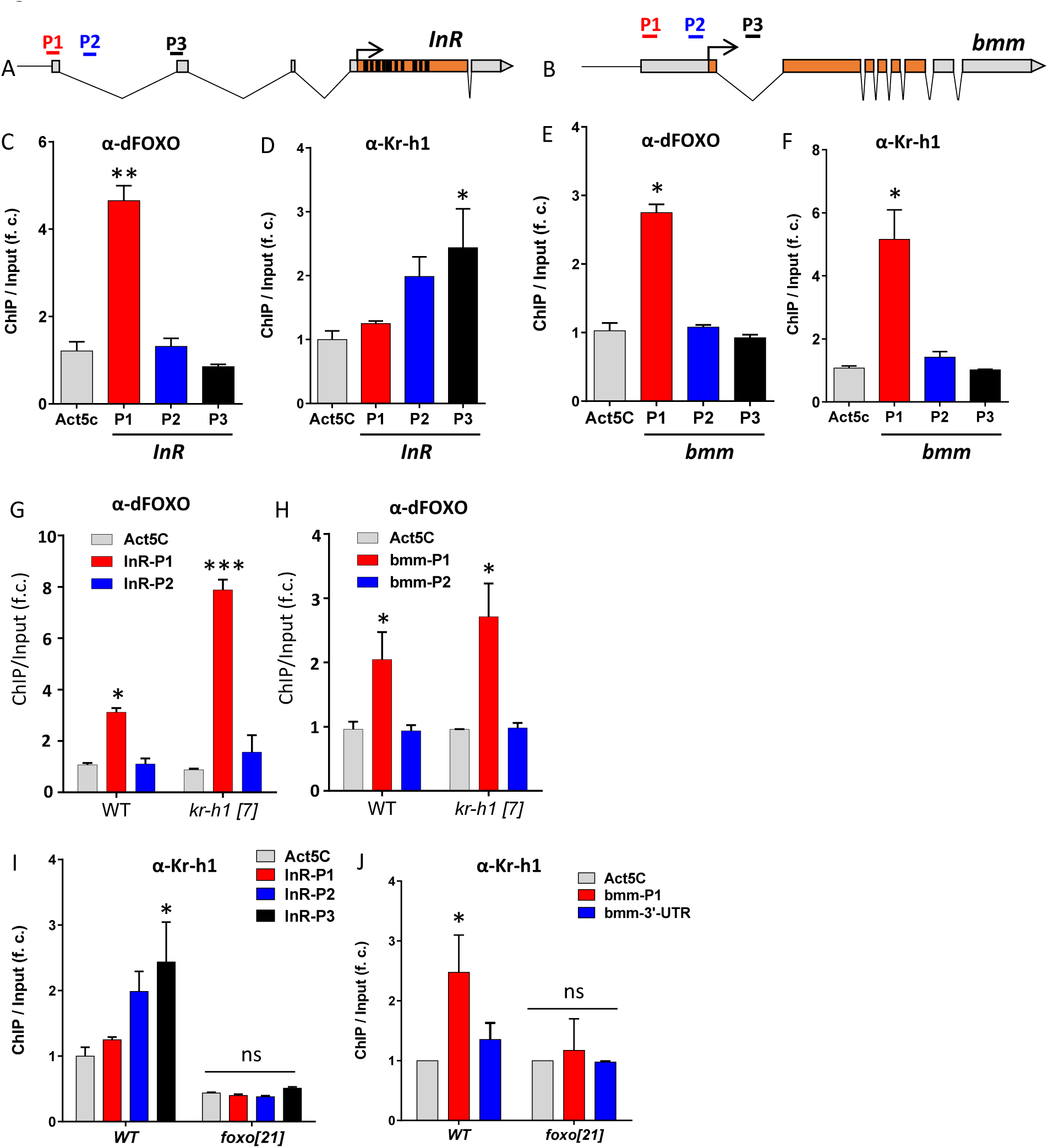
Kr-h1 binds to the promoter of brummer lipase and insulin receptor adjacent to dFOXO binding sites. (A). Schematic graph shows insulin receptor (*InR*) locus. P1 region contains a canonical FOXO binding motif (GTAAATAA), while putative mammalian Kruppel binding sits are found in all three regions (based on motif search on the Jaspar database, jaspar.genereg.net). (B). Schematic graph shows brummer lipase (*bmm*) locus. P1, P2 and P3 are corresponding to the target sites tested in ChIP-PCR analysis. P1 region contains a canonical FOXO binding motif, while putative mammalian Kruppel binding sits are found in all three regions. (C). ChIP-PCR analysis on dFOXO binding to *InR* promoter. (D). ChIP-PCR analysis on Kr-h1 binding to *InR* promoter.Each bar represents mean ± SE of three biological replicates. Statistical significance is assessed by one-way ANOVA. (E). ChIP-PCR analysis on dFOXO binding to *bmm* promoter. (F). ChIP-PCR analysis on Kr-h1 binding to *bmm* promoter. (G). dFOXO binding to *InR* promoter (P1 region) is enhanced in fasted *Kr-h1* mutants. Interaction is statistically significant, *p*<0.0001. (H). dFOXO binding to *bmm* promoter (P1 region) is slight enhanced fasted *Kr-h1* mutants. Interaction is not statistically significant, *p*=0.5862. (I). Kr-h1 binding to InR promoter is abolished in fasted *dfoxo[21]* mutants. (J). Kr-h1 binding to bmm promoter is abolished in fasted *dfoxo[21]* mutants. Each bar represents mean ± SE of three biological replicates. Statistical significance is assessed by one-way ANOVA, followed by Dunnett’s multiple comparisons (* p<0.05, ns: not significant).

Binding of dFOXO and Kr-h1 to these putative sites was determined by ChIP-PCR analysis in fasted animals. At *InR*, dFOXO binding was strongest in the P1 region located at the 5’-UTR region (Fig. 4C), while Kr-h1 bound most strongly to the P3 regions (Fig. 4D). At *bmm* lipase, both dFOXO and Kr-h1 bound with highest affinity in the P1 region (Fig. 4E & 4F). The co-localization of Kr-h1 and dFOXO binding suggests these factors could interact at promoters to control the transcriptional activation of the key metabolic genes, and *bmm* lipase in particular.

### Kr-h1 represses dFOXO binding to the promoter of *InR* and *bmm*

Kr-h1 may repress dFOXO activity by inhibiting its binding at response elements in *bmm* and *InR*. We performed a ChIP-PCR to test this possibility using anti-dFOXO antibody and *Kr-h1[7]* mutants. dFOXO binding to the *InR* P1 region was increased from 2.9-fold relative to negative control (Act5C) in fasted wildtype to 8.95-fold in fasted *Kr-h1* mutants (Two-way ANOVA, interaction p<0.0001) (Fig. 4G). In contrast, dFOXO binding to the *bmm* P1 region was slightly but non-significantly increased from 2.1-fold in fasted wildtype to 2.8-fold in fasted *Kr-h1* mutants (Two-way ANOVA, interaction *p*=0.5862) (Fig. 4H). At the *InR* promoter in particular, inhibition of dFOXO-DNA interaction may be one mechanism by which Kr-h1 modulates dFOXO transcriptional activity. Notably, in a reciprocal experiment with anti-Kr-h1 antibody, the binding of Kr-h1 to *InR* and *bmm* promoters was abolished in *dfoxo[21]* mutants (Fig. 4I & 4J). These data suggest that Kr-h1 may be recruited after dFOXO binds to the promoters of target genes, and Kr-h1 subsequently modulates the transcriptional activities of dFOXO through interfering with dFOXO-DNA interactions.

### Kr-h1 expresses in adipose tissue to control larval development and lipid metabolism

To determine where Kr-h1 and dFOXO interact *in vivo*, we first examined the tissue-specific expression of Kr-h1 using our anti-Kr-h1 antibodies. Interestingly, Kr-h1 expressed in larval ring gland, especially in corpora allata (CA), the production sites of JH (Fig. 5A). No expression of Kr-h1 was detected in insulin producing cells (IPCs) (Fig. 5B). Kr-h1 expressed highly in body wall muscle and midgut muscle (Fig. 5C), and in fasted fat body (Fig. 5D). Upon fasting, dFOXO is activated by de-phosphorylation and subsequent nuclear translocation ^37^.

**Fig 5.**
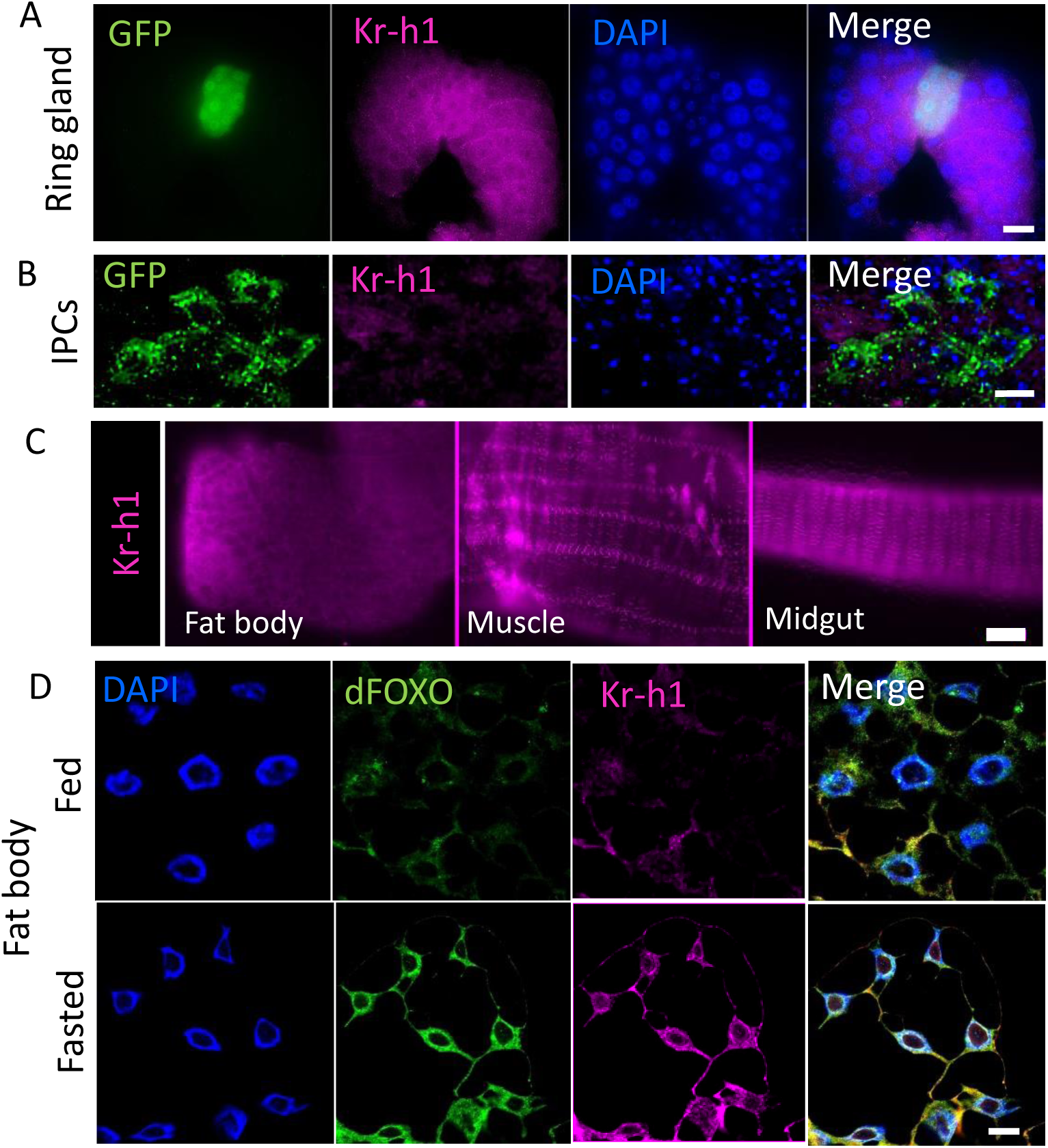
Tissue-specific expression pattern of Kr-h1. (A). Kr-h1 expressed in ring gland of Aug21-gal4>UAS-GFP.nls larvae. CA is labeled by GFP staining. Scale bar: 20 μm. (B). Kr-h1 does not express in IPCs of dilp2-gal4>UAS-mCD8::GFP larvae. A cluster of IPCs is labeled by GFP staining. Scale bar: 10 μm. (C). Kr-h1 expressed in larval body wall muscle and midgut muscle. Scale bar: 20 μm. (D). Nuclear co-localization of Kr-h1 and dFOXO in fat body upon fasting. Larvae at 90 hr AEL were fasted for 16 hr in culture vials with wet kimwipe soaked with PBS. Fat body cells were dissected and staining with anti-Kr-h1 and anti-dFOXO antibodies.Scale bar: 10 μm.

Nuclear translocation of both dFOXO and Kr-h1 was increased in fasted larval fat body (Fig. 5D). Thus, dFOXO and Kr-h1 may interact in fat body at the genome to co-regulate the transcriptional activation of target genes.

dFOXO expressed in fat body regulates lipid metabolism and Dilp2 production from brain IPCs^38^. To determine from which tissue Kr-h1 regulates lipid metabolism and larval development, we knocked down *Kr-h1* message through RNA interference (RNAi) with specific Gal4 drivers. Knockdown of *Kr-h1* in fat body (r4-gal4) and muscle (Mhc-gal4) delayed the pupariation, while knockdown in gut, IPCs and CA showed no effects on larval development (Fig. 6A). Since fat body is the major site for triglyceride storage in *Drosophila*, we further examined the role of Kr-h1 in the regulation of lipid metabolism in fat body. Consistently, fat body-specific knockdown of *Kr-h1* induced *bmm* transcription, while overexpression of *Kr-h1* repressed it (Fig. 6B). Fat body-expressed *Kr-h1* also increased TAG levels (Fig. 6C). Thus, adipose-expressed Kr-h1 is essential for larval development and metabolic regulation.

**Fig 6.**
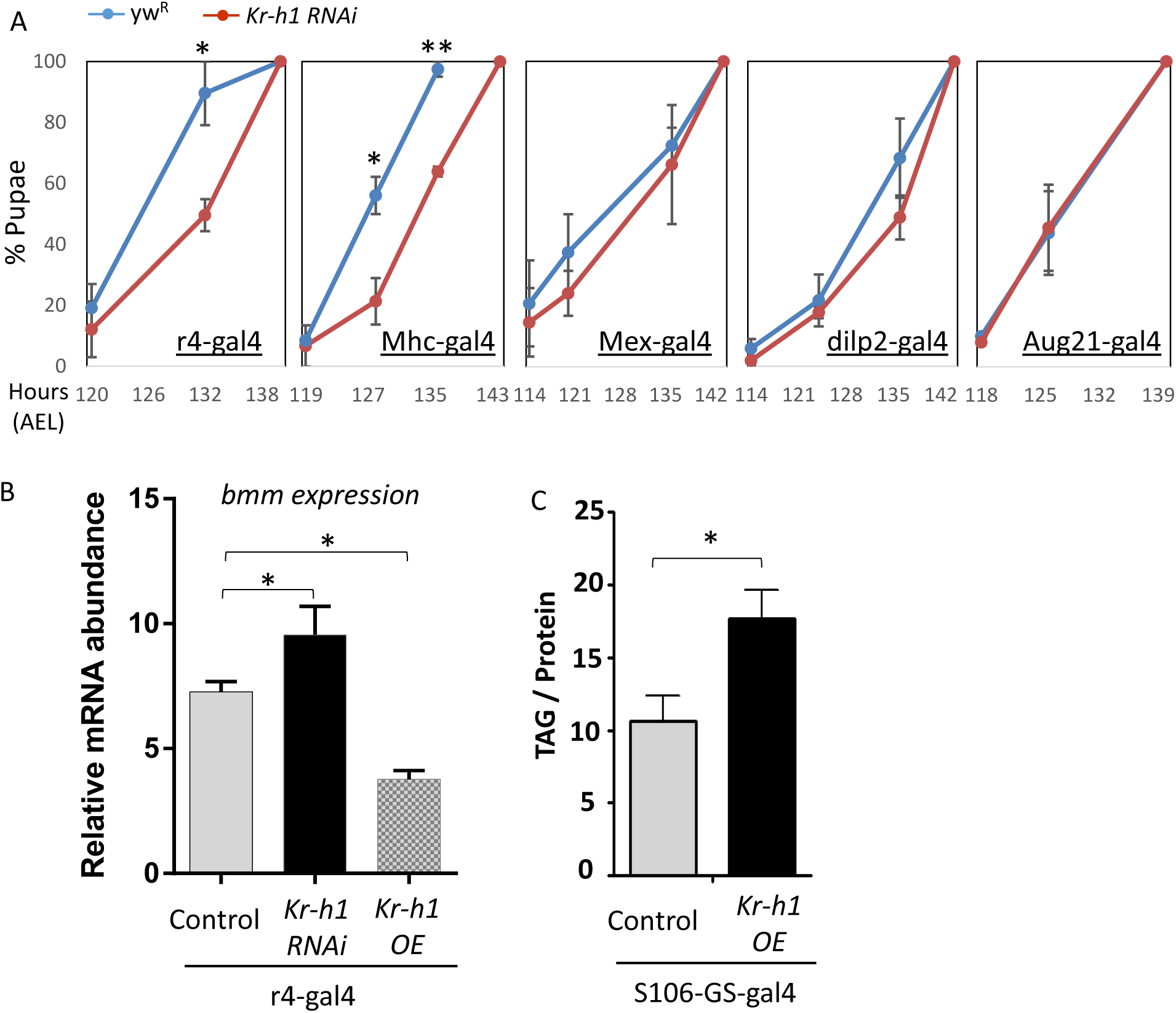
Fat body-expressed Kr-h1 regulates larval development and lipid metabolism. (A). Knockdown of *Kr-h1* expression in fat body (r4-gal4) and muscle (Mhc-gal4) delayed the pupariation. Knockdown of *Kr-h1* in gut (Mex-gal4), IPCs (dilp2-gal4) and CA (Aug21-gal4) shows no effects on larval development. The Kr-h1 RNAi line was backcrossed into a *yw*^*R*^ background for five generations prior to developmental timing experiments. Data are represented as mean ± SE of three trials. Student t-test (** p<0.01, * p<0.05) (B). Fat body-specific knockdown of *Kr-h1* induced *bmm* transcription, while overexpression of *Kr-h1* in fat body repressed it. (C). Fat body-specific overexpression of *Kr-h1* increased TAG levels. Data are represented as mean ± SE of three trials. Student t-test or one-way ANOVA (* p<0.05).

### Juvenile hormone signaling regulates lipase *bmm* through dFOXO

The interaction between Kr-h1 and dFOXO has the potential to integrate development and nutrient signaling. Nutrient signaling through FOXO involves insulin, AMPK, SIRT and JNK in both insect and mammals alike ^6, 13^. On the other hand, the upstream regulators of Kruppel-like factors are poorly characterized in vertebrates, but among insects Kr-h1 is decisively regulated by JH, a key hormonal signal involved in molting and metamorphosis ^20^. In particular, JH induces the transcription of *Kr-h1* via the JH receptor Methoprene-tolerant (Met) ^21, 39^. In this capacity, recent studies suggest that JH and Met are involved in not only development programming, but also in metabolic control ^9, 40-42^, although how JH affects metabolism is fundamentally unknown.

Given that Kr-h1 and dFOXO functionally interact to control lipid metabolism, we examined if this feature provides a way for JH to affect metabolic regulation through *bmm* transcription. Consistent with previous studies ^9^, triglyceride levels were reduced in flies where the corpora allata were genetically removed (CAX) (Fig. 7A). Conversely, wild-type flies exposed to the JH analog (JHA) methoprene had elevated TAG contents compared to controls (Fig. 7B). Additionally, *Met* mutations had also down-regulated TAG levels (Fig. 7C) and up-regulated *bmm* mRNA (Fig. 7D). Met also genetically interacts with dFOXO to regulate the mRNA expression of *bmm* (Fig. 7D). JH may therefore regulate *bmm* via interaction with dFOXO. Supporting this prediction, methoprene treatment inhibited the expression of *bmm* in wildtype flies, but not in *dfoxo[21]* mutants (Fig. 7E). Furthermore, fasting reduced JH titers about 2-fold in both female and male flies (Fig. 7F). But while it is known that JH positively regulates Kr-h1 transcription^20, 22^, *Kr-h1* mRNA did not change upon fasting (Fig. 7G). On the other hand, methoprene treatment was sufficient to induce *Kr-h1* transcription (Fig. 7H). Overall these results indicate that JH signaling interacts with dFOXO to regulate lipid metabolism and lipase gene expression, but the specific role of Kr-h1 in this process remains to be elucidated.

**Fig 7.**
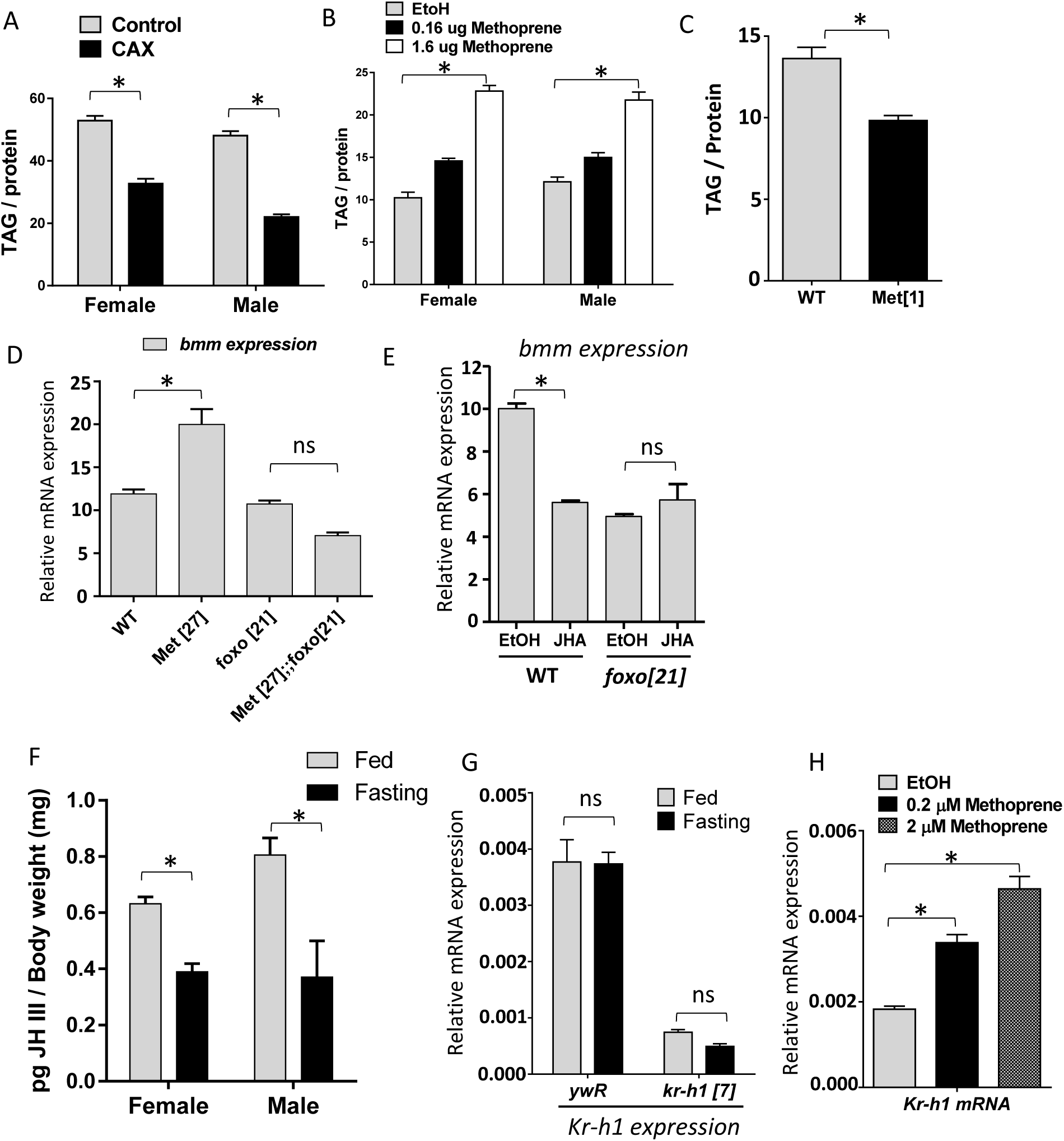
Juvenile hormone signaling regulates TAG lipase *bmm* through dFOXO. (A). TAG levels are reduced in CA ablation (CAX) flies. Each bar represents mean ± SE of three biological replicates. Student t-test (* p<0.05). (B) Flies exposed to JH analog (JHA) methoprene show increased TAG levels. Each bar represents mean ± SE of three biological replicates. One-way ANOVA (* p<0.05). (C). *Met* mutants have reduced TAG levels. Student t-test (* p<0.05). (D). Genetic interaction between *Met* and *dfoxo* in the regulation of *bmm* transcripts. *Bmm* transcription is up-regulated in *Met* mutants, which was rescued by *dfoxo*^*21*^ mutants. One-way ANOVA (* p<0.05, ns: not significant). (E). JH analog (JHA) methoprene treatment led to reduced *bmm* expression in wildtype female flies, but not in *dfoxo*^*21*^ mutant flies. Each bar represents mean ± SE of three biological replicates. Statistical significance is assessed by two -way ANOVA (* p<0.05, ns: not significant) (F). JH titer is decreased upon fasting. 10-day-old adult flies were fasted (in culture vial with wet kimwipe soaked with PBS) for 16 hours before collected for JH quantification. Each bar represents mean ± SE of 5∼7 biological replicates. Statistical significance is assessed by student t-test (* p<0.05). (G). The mRNA expression of Kr-h1 did not change upon fasting. (H). Methoprene treatment induced *Kr-h1* transcription. Each bar represents mean ± SE of three biological replicates. One -way ANOVA (* p<0.05, ns: not significant).

## Discussion

Transcriptional coordination is a key process contributing to metabolic homeostasis ^43^. Multiple transcription factors interact at their genomic binding sites to enhance transcriptional specificity and pleiotropic functions of metabolic pathways. As a key node in the metabolic network, forkhead transcription factor FOXO has been shown to interact with diverse transcription co-factors and thereby integrate signals to control metabolism and oxidative stress ^6, 13^. Intriguingly, in recent genomic studies ^44-46^, the enriched FOXO binding at specific genes does not always correlate to elevated transcriptional output, suggesting there exists inhibitory or inertial mechanisms to repress FOXO when it is already bound to target genes.

Here we find that *Drosophila* Kruppel-like factor Kr-h1 acts as a repressor of dFOXO to modulate induction of two dFOXO target genes, *InR* and *bmm*. Like other FOXO interacting partners, Kr-h1 physically binds to dFOXO and inhibits the expression of dFOXO targets by influencing the binding affinity of dFOXO to DNA. The transcriptional activity of FOXO is typically regulated in two layers. The first and probably most important regulation is through PTM, including phosphorylation, acetylation and ubiquitination ^6^. PTM of FOXO proteins can affect its subcellular localization (by phosphorylation), DNA binding affinity (by acetylation) and protein degradation (by ubiquitination). Interestingly, the effects of acetylation on FOXO factors seem to be quite different from those by acetylation on KLFs. Acetylation of FOXO by co-factor CBP/p300 weakens the FOXO binding to its DNA targets ^47^, while CBP/p300 acetylated KLF1 shows increased transcriptional activation of target gene beta-globin ^29^. The second mechanism for the regulation of FOXO activity is through the interaction between FOXO and other transcription factors or co-factors. FOXO factors have been shown to interact with diverse transcription factors (e.g. Smad3/4, PGC-1α, STAT3) that often potentiate the expression of FOXO target genes ^13^. Kr-h1 identified in our study presents another example for this type of modulatory regulation, although the interaction between Kr-h1 and dFOXO results in transcription repression, instead of activation.

While we do not fully resolve how Kr-h1 blocks dFOXO activity, it seems that Kr-h1 can inhibit dFOXO binding to its DNA targets. This result is similar to previous studies showing reduced FOXO-DNA binding upon interaction with androgen receptor (AR) ^48^ and with peroxisome proliferator-activated receptor-γ (PPARγ) ^49^. Alternatively, Kr-h1 may act by inhibiting recruitment of dFOXO-coactivators (e.g. SIRT or CBP/p300) or by sequestering these coactivators away from dFOXO. Kr-h1 may also recruit additional co-repressors (e.g. CtBP or Sin3-HDAC) to the dFOXO transactivation sites to block the transcriptional activation of target genes. The N-terminal Q-rich domain of KLFs is crucial for the recruitment of co-repressors CtBP and Sin3A ^50^. In our co-immunoprecipitation assays, the Q-rich domain of Kr-h1 strongly binds to dFOXO, suggesting Kr-h1 might inhibit dFOXO activity through recruiting co-repressors. One possible candidate is an Sds3-like gene (CG14220), which was previously found to co-immunoprecipitate with Kr-h1 ^51^. Sds3-like gene family proteins can form co-repressor complex with Sin3A and HDAC to inhibit gene transcription via interactions with sequence-specific transcription factors ^52^.

KLFs have well documented roles in cell proliferation, differentiation and apoptosis ^50^. A function for KLFs in lipid metabolism and insulin signaling complements recent studies where KLFs function in cellular metabolic regulation, such as gluconeogenesis ^53, 54^. Likewise, KLF15 deletion in mice produces hypoglycemia and impaired amino acid catabolism upon fasting ^54^.Additionally, KLF5 heterozygous mice are resistant to high-fat diet-induced obesity. SUMOylation modulates the transcriptional activities of KLF5 and its association with peroxisome proliferator-activated receptor-delta (PPAR-delta) to control the expression of carnitine-palmitoyl transferase-1b (Cpt1b), uncoupling proteins 2 and 3 ^55^. Interestingly, KLF4 has recently been identified as a direct target gene of FOXO-mediated transcription during B cell development ^56^, suggesting a potential interaction between KLF and FOXO transcriptional regulatory network.

Our ChIP-PCR studies suggest that *Drosophila* KLF Kr-h1 transcriptionally controls many metabolic genes, including some of the key dFOXO targets (e.g. *InR* and triglyceride lipase *bmm*). While it is not known whether *Drosophila* Kr-h1 could broadly interact with dFOXO across the genome, such a genome-wide interaction between mammalian FOXO factors and KLFs has been suggested by a recent meta-analysis ^57^. Because Kr-h1 plays an important role in morphogenesis during *Drosophila* development ^20^, the interplay between Kr-h1 and dFOXO raises the possibility that Kr-h1 coordinates growth and development through insulin/dFOXO-mediated metabolic regulation.

In insects, Kr-h1 is one of the key effectors of JH signaling, an important hormonal pathway governing insect molting, metamorphosis and reproduction. Recent studies reveal that JH also participates in the regulation of carbohydrate and lipid metabolism ^9, 40-42, 58^. In Tsetse flies, JH and insulin co-regulate the expression of TAG lipase and inhibit lipolysis ^9^, which is similar to our observation that TAG metabolism and lipase *brummer* expression are regulated by JH and its receptor Met. JH signaling genetically interacts with insulin/dFOXO to control larval growth rate and define final body size ^10^. Thus, the transcriptional co-regulation of lipid metabolism by *Drosophila* Kr-h1 and dFOXO may contribute to a novel mechanism through which JH interacts with insulin signaling to integrate metabolism and growth during larval development.

Since both Kr-h1 and dFOXO express highly in metabolic tissues (fat body and muscle) of *Drosophila*, it is likely that the two transcription factors co-regulate many key metabolic genes in these tissues. On the other hand, these metabolic tissues also contribute significantly to other insect physiology and organismal functions, such as stress resistance and aging that are tightly regulated by insulin/dFOXO signaling ^59^ and linked to JH signaling ^40^. Therefore, the interplay between Kr-h1 and dFOXO may contribute to the regulation of these adult physiological processes. Identifying the key co-factors and downstream events of Kr-h1/dFOXO transcriptional network may advance our understanding of the integrated regulation by JH and insulin signaling of metabolic, developmental and aging pathways.

## Materials and Methods

### Fly Husbandry and Stocks

Flies were maintained at 25°C, 40% relative humidity and 12 -hour light/dark. Adults were reared on agar-based diet with 0.8% cornmeal, 10% sugar, and 2.5% yeast (unless otherwise noted). Fly stocks used in the present study are: *Kr-h1[7]* or *Kr-h1 [k04411]* ^19, 30^ (Bloomington # 10381, backcrossed to *yw*^*R*^), *Kr-h1* RNAi lines (Bloomington # 50685, VDRC #107935), *Kr-h1* EP line #EP2289 ^23, 60^, UAS-Kr-h1-LacZ ^24^, *foxo[21]* ^61^, *Met[1]* ^62^, *Met[27]* ^62^, r4-gal4 (Bloomington # 33832), Mhc-gal4 ^63^, Mex-gal4 ^64^, dilp2-gal4 ^65^, Aug21-gal4 ^66^, S106-GS-gal4 ^67^, UAS-GFP.nls (Bloomington # 4775), UAS-mCD8::GFP (Bloomington # 5137). Double mutants were made by crossing *Kr-h1[7]* or *Met[27]* to *foxo[21]* respectively. Corpus allatum (CA) ablation flies (named CAX flies) are generated in our laboratory as previously described ^40^. *yw*^*R*^ flies were used a wild-type flies in most of the experiments. For methoprene treatment, adult flies were exposed for 24∼48 hours to various concentrations of methoprene applied to the side of culture vials.

### Kr-h1 Antibody and Western Blot

Kr-h1 polyclonal antibody was generated in rabbits against the short peptide sequence ‘LIEHFKRGDLARHG’ (Covance, Dedham, MA, USA) and affinity purified (Thermo Fisher Scientific, Waltham, MA, USA). The antibody recognized three major bands in western blots (Fig. 1C). These bands may be corresponding to the three isoforms of Kr-h1 (α, β, γ). All western blots were performed per the following procedures: Fly tissues or cells were homogenized in RIPA buffer (Thermo Fisher Scientific, Waltham, MA, USA) with protease inhibitors (Sigma-Aldrich, St Louis, MO, USA). Supernatant was incubated with NuPAGE LDS loading buffer (Thermo Fisher Scientific, Waltham, MA, USA) at 70 ^0^C for 10 min. About 20 μg of denatured protein was separated on 4∼12% Bis-Tris precast gels (Thermo Fisher Scientific, Waltham, MA, USA) and transferred to PVDF membranes. Following incubation with primary and secondary antibodies, the blots were visualized with Pierce ECL Western Blotting Substrate (Thermo Fisher Scientific, Waltham, MA, USA). Other antibodies used in the present study are Phospho-*Drosophila* Akt antibody (Ser505) (#4054S, Cell Signaling Technology, Danvers, MA, USA), Akt antibody (#9272S, Cell Signaling Technology).

### Quantitative RT–PCR

Total RNA was extracted using Trizol reagent (Thermo Fisher Scientific, Waltham, MA, USA) from 10 ∼15 synchronously staged larvae or whole adult flies. DNase-treated total RNA was quantified and about 500 ng of total RNA was reverse transcribed to cDNA using iScript cDNA Synthesis Kit (Bio-Rad, Hercules, CA, USA). QPCR was performed with an ABI prism 7300 Sequence Detection System (Thermo Fisher Scientific, Waltham, MA, USA). Three to five biological replicates were used for each experimental treatment. mRNA abundance of each gene was normalized to the expression of ribosomal protein L32 (*RpL32* or *rp49*) by the method of comparative CT . Primer sequences are listed in Supplementary Table S1.

### Pupariation timing analysis

Synchronized eggs were placed on 35 x 10 mm petri dishes containing standard medium (see above) at 20∼30 eggs per dish. The numbers of pupae were recorded 2∼3 times every day around 120 hours AEL till all larvae molt into pupae.

### Metabolic assays

All metabolic analyses were performed as previously described ^67, 68^. For TAG assay, 25 staged larvae or six adult flies were collected and homogenized in 1xPBS containing 0.1% Tween 20 and TAG was quantified using Thermo Scientific(tm) Triglycerides Reagent (Thermo Fisher Scientific, Waltham, MA, USA). For glycogen measurement, samples were digested with amyloglucosidase (Sigma-Aldrich, St Louis, MO, USA) and glucose contents were quantified using Thermo Scientific(tm) Glucose Hexokinase Reagents (Thermo Fisher Scientific, Waltham, MA, USA).The relative level of each metabolite was obtained by normalizing the metabolites to total protein.

### Immunoprecipitation and pull-down

All the immunoprecipitation and pull-down experiments were conducted in *Drosophila* Kc167 cells adapted to serum-free culture medium (*Drosophila* Schneider Medium). Either full-length (Kr-h1 α-isoform) or partial gene products were cloned into *Drosophila* Gateway Vectors with N-terminal tags (FLAG and HA) following *Drosophila* Gateway Vectors protocols (https://emb.carnegiescience.edu/*<u>Drosophila</u>*<u>-gateway-vector-collection</u>). About 1 μg of constructs were transfected to 2 x 10^6^ Kc167 cells using Effectene reagent (Qiagen, Hilden, Germany). Two days after transfection, cells were harvested and lysed in NP-40 lysis buffer (Thermo Fisher Scientific, Waltham, MA, USA) with proteinase inhibitors (Sigma-Aldrich, St Louis, MO, USA). To pull-down target proteins, total protein extracts were incubated with proper antibodies and Dynabeads Protein A (Thermo Fisher Scientific, Waltham, MA, USA). Following pull-down, western blotting was performed to examine protein complex. Antibodies used in pull-down and western blots include rabbit anti-Kr-h1 and anti-dFOXO produced in our laboratory, rabbit anti-HA (Covance, Dedham, MA, USA), and mouse anti-FLAG (Sigma-Aldrich, St Louis, MO, USA). Nuclear extracts for immunoprecipitation were conducted with a nuclear extraction kit (Active motif, Carlsbad, CA, USA).

### Immunohistochemistry and imaging

To examine the tissue-specific expression of Kr-h1 and its co-localization with dFOXO, various larval tissues were dissected from fed or fasted 3^rd^ instar larvae (90 hr AEL) (For fasting, larvae were placed onto wet kimwipe soaked with 1 x PBS for 16 hours). Tissue immunostaining were performed as previously described ^46^, using slowFade mounting solution with DAPI (Thermo Fisher Scientific, Waltham, MA, USA). Samples were imaged with a Zeiss 510 laser scanning confocal microscope or an Olympus BX51WI upright epifluorescence microscope equipped with Hamamatsu Flash 4.0 Plus CMOS Camera. Antibodies used in immunohistochemistry included: rabbit anti-Kr-h1 (1:200) (this study), anti-dFOXO (1:200) ^69^, anti-GFP (Sigma-Aldrich, St Louis, MO, USA), anti-rabbit IgG-DyLight 488 (1:300) anti-rabbit IgG-Alexa Fluor 594 (1:300) and anti-Guinea pig IgG-DyLight 488 (1:300) (Jackson ImmunoResearch, West Grove, PA, USA).

### Chromatin immunoprecipitation (ChIP)

ChIP was conducted as previously described ^46^. About 50 staged larvae were used in each sample. Flies were homogenized and cross-linked in 1xPBS containing 1% formaldehyde. The fly nuclear extractions were sonicated using a Branson 450 sonicator to break down the chromatins. Immunoprecipitation was performed using Dynabeads Protein A and anti-Kr-h1 and anti-dFOXO antibodies. Following the wash with LiCl and TE buffer, the DNA-protein complex was eluted, reverse cross-linked, digested with Proteinase K and RNase. Kr-h1-bound or dFOXO-bound DNA fragments were purified and used as templates in qPCR analysis. Binding enrichment was calculated as the fold change between ChIP DNA vs. input DNA (Chromatin extracts before immunoprecipitation). The binding to the coding region of Actin (*Act5C*) was used as negative controls.

### Juvenile hormone quantification

For each sample, 197-200 individual flies (7∼10-day-old) were placed in 500 μl hexane in a glass vial with a Teflon cap insert and stored at −80°C prior to analysis. To extract the hormone, the flies were crushed with a Teflon tissue grinder. The resultant homogenate was centrifuged at 3500 rpm for 5 min, and the supernatant was removed to clean vial. Extraction was conducted three times, combining the resultant supernatant from each sample. The gas chromatography/mass spectrometry (GC–MS) method ^70^, as modi?ed ^71, 72^, was used to quantify juvenile hormone (JH). Samples were eluted through aluminum oxide columns successively with hexane, 10% ethyl ether–hexane and 30% ethyl ether–hexane. Samples were subjected to a second series of aluminum oxide elutions (30% ethyl ether-hexane then 50% ethyl-acetate– hexane) after derivatization with methyl-d alcohol (Sigma-Aldrich, St Louis, MO, USA) and trifluoroacetic acid (Sigma-Aldrich, St Louis, MO, USA). Purified samples were analyzed on an HP 7890A Series GC (Agilent Technologies, Santa Clara, CA, USA) equipped with a 30 m x 0.25 mm Zebron ZB-WAX column (Phenomenex, Torrence, CA, USA) and coupled to an HP 5975C inert mass selective detector with helium as the carrier gas. MS analysis occurred in the SIM mode, monitoring at m/z 76 and 225 to ensure specificity for the d3-methoxyhydrin derivative of JH III. Total abundance was quantified against a standard curve of derivatized JH III and using farnesol (Sigma-Aldrich, St Louis, MO, USA) as an internal standard. The detection limit is approximately 1 pg.

### Statistical analysis

GraphPad Prism 6 (GraphPad Software, La Jolla, CA) was used for statistical analysis. To compare the mean value of treatment groups versus that of control, either student t-test or one-way ANOVA was performed using Dunnett’s test for multiple comparison. The effects of mutants on starvation responses was analyzed by two-way ANOVA, including Tukey multiple comparisons test.

## Acknowledgements

We thank Bloomington *Drosophila* Stock Center, *Drosophila* Genomics Resource Center, and Vienna Drosophila Resource Center for fly stocks and cDNA clones. We thank Drs. Yannick Beck, Heather Broihier, Fabio Demontis, Tzumin Lee, Franck Pichaud, Geoff Richards, Eric Rulifson, Graham H. Thomas, Thomas Wilson for providing fly stocks and reagents. This work was supported by National Institutes of Health/National Institute on Aging grant R37 AG024360 to MT, R00 AG048016 to H.B.

Mention of trade names or commercial products in this publication is solely for the purpose of providing specific information and does not imply recommendation or endorsement by the U.S. Department of Agriculture. USDA is an equal opportunity provider and employer.

## Author Contributions

Conceived and designed the experiments: MT HB. Performed the experiments: PK KC YL MB GK RT WZ SP CSB HB. Analyzed the data: PK KC CSB MT HB. Wrote the paper: CSB SL MT HB. All authors reviewed and approved the final version of this manuscript.

## Additional Information

**Competing Interests:** The authors declare that they have no competing interests.

## Supplementary Information

Supplementary Table S1. Putative KLF binding sites in the promoters of *InR* and *bmm*.

Supplementary Table S2. List of Primers

Supplementary figure S1. Original full-length blots for Figure 1C

Supplementary figure S2. Original full-length blots for Figure 2A

Supplementary figure S3. Original full-length blots for Figure 3A

Supplementary figure S4. Original full-length blots for Figure 3B

Supplementary figure S5. Original full-length blots for Figure 3C

**Supplementary Table S2.**
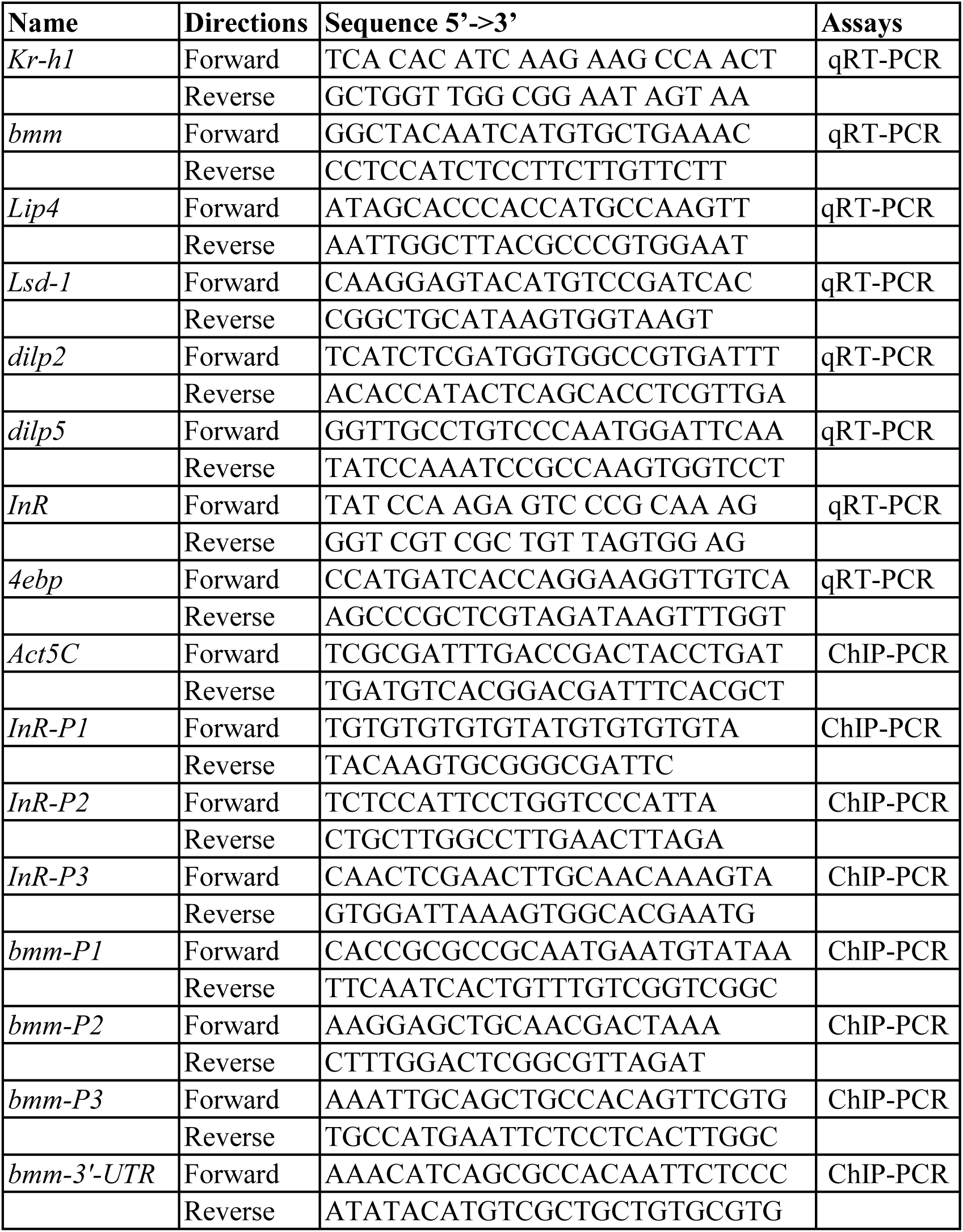
List of Primers.

**Supplementary Table S1.**
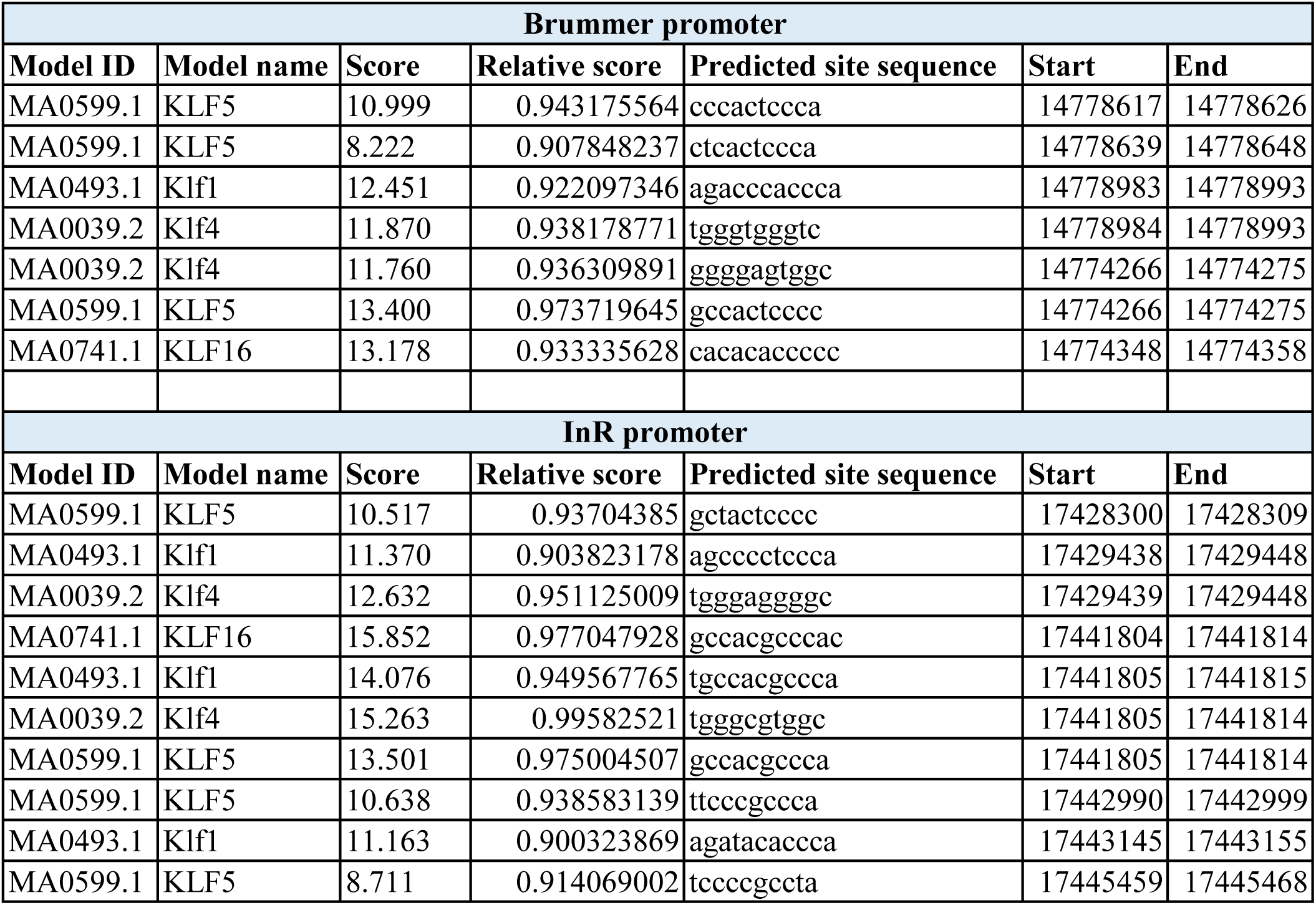
Putative KLF binding sites in the promoters of InR and bmm.

**Supplementary figure S1.**
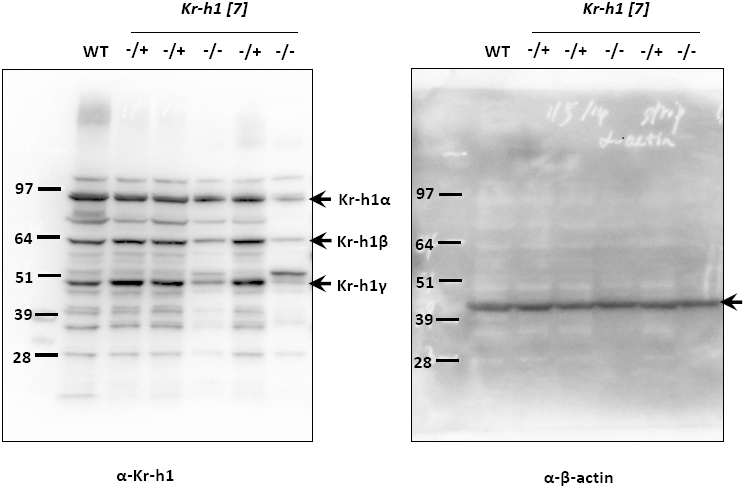
Original full-length blots for Figure 1C.

**Supplementary figure S2.**
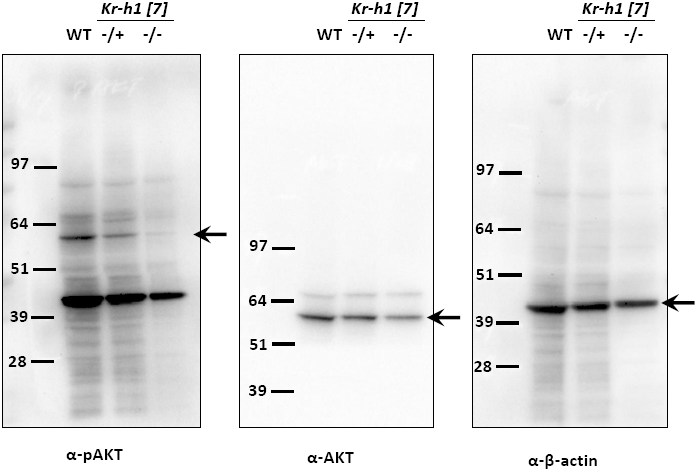
Original full-length blots for Figure 2A.

**Supplementary figure S3.**
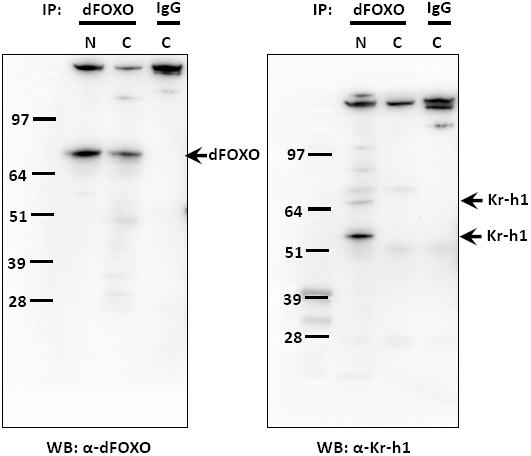
Original full-length blots for Figure 3A.

**Supplementary figure S4.**
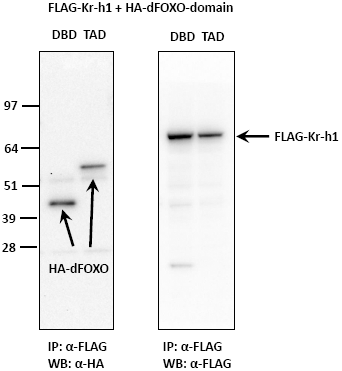
Original full-length blots for Figure 3B.

**Supplementary figure S5.**
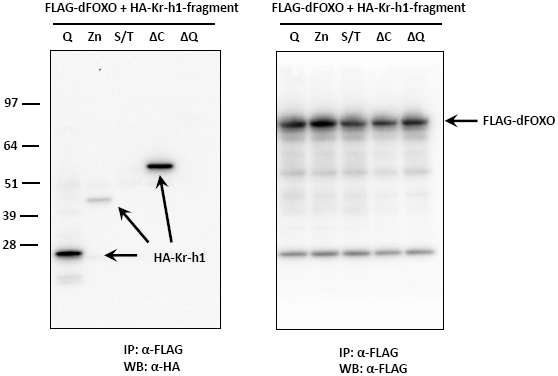
Original full-length blots for Figure 3C.

